# Sea Turtle Facial Recognition Using Map Graphs of Scales

**DOI:** 10.1101/2021.06.03.446936

**Authors:** Karun K. Rao, Lars C. Grabow, Juan Pablo Muñoz-Pérez, Daniela Alarcón-Ruales, Ricardo B. R. Azevedo

**Affiliations:** Department of Chemical and Biomolecular Engineering, University of Houston, Houston, TX 77204-4004, USA; Texas Center for Superconductivity at the University of Houston, Houston, TX 77204-5002, USA; UNC-Chapel Hill & Universidad San Francisco de Quito (USFQ) Galápagos Science Center (GSC) Av. Alsacio Northia, Isla San Cristobal, Galápagos, Ecuador; Faculty of Science and Engineering, University of the Sunshine Coast, QLD, Australia; Department of Biology and Biochemistry, University of Houston, Houston, TX 77204-5001, USA

**Keywords:** Animal Biometrics, Coherent Point Drift, Facial Recognition, Individual Identification, Map Graphs, Networks, Sea Turtles

## Abstract

1. Individual identification of sea turtles is important to study their biology and aide in conservation efforts. Traditional methods for identifying sea turtles that rely on physical or GPS tags can be expensive, and difficult to implement. Alternatively, the scale structure on the side of a turtle’s head has been shown to be specific to the individual and stable over its lifetime, and therefore can be used as the individual’s “fingerprint”.
2. Here we propose a novel facial recognition method where an image of a sea turtle is converted into a graph (network) with nodes representing scales, and edges connecting two scales that share a border. The topology of the graph is used to differentiate species.
3. We additionally develop a robust metric to compare turtles based on a correspondence between nodes generated by a coherent point drift algorithm and computing a graph edit distance to identify individual turtles with over 94% accuracy.
4. By representing the special and topological features of sea turtle scales as a graph, we perform more accurate individual identification which is robust under different imaging conditions and may be adapted for a wider number of species.

## 1. Introduction

Computer-assisted individual photo identification has been employed in a variety of species, including whale sharks (Arzoumanian et al. 2005), seadragons (Martin-Smith, 2011), salamanders (Gamble et al, 2008), leopards (Miththpala, et al. 1989), and sea turtles (Gatto et al., 2018; Jean et al., 2010; Reisser et al., 2008). Visually comparing all possible combinations of individuals is impractical as the number of images grows, so an automated method for performing individual identification is used, typically based on identifying biological markers of the specific species. These natural markers are often specific patterns of spots, lines, or unique markings/patterns which can be automatically detected based on contrast enhancement or line/edge detection filters. In many cases, such as in sea turtles, the spatial distribution (position and number) of the markers is not enough to uniquely identify an individual.

A significant challenge with marine animal imaging is that images are not generated in a consistent or controlled manner; images are collected under different conditions of light, visibility, and background, at different distances, from different angles, with different cameras, and by different observers. Existing software for performing individual recognition of sea turtles and other marine animals relies mainly on manual region selection and automated feature identification on the plastron of the turtle using the features generated by SIFT (Kisku et al. 2007), SURF (Leonardis et al. 2006), or ORB (Rublee et al. 2011). The main limitation of this approach is the reliance on the SIFT-based descriptor (Beugeling & Branzan-Albu, 2014), which has little physical relevance to the turtles’ scale structures and often relies on detection of general edges or regions of intensity that could vary widely based on the region analyzed. Furthermore, this approach produces inconsistent results under the wide variety of water and lighting conditions in which turtles are photographed, without extensive preprocessing (Calmanovici et al. 2018). Additional false negatives are generated when the tilt of the turtles’ head differs by more than ∼30% which is difficult to control when taking pictures of moving turtles under water (den Hartog & Reijns, 2019). Once identified, features can be compared across databases in many different ways using a variety of hierarchical methods to rank the most similar individuals (Duyck et al., 2015).

Reisser et al. 2008 explored descriptors of the turtles’ facial scale structures that were more representative physically using a purely qualitative visual inspection of the scale pattern to validate tagging data, and identify individuals. Recently, Jean et al. 2010 assigned each facial scale a unique three digit ID number encoding the relative position of the scale with the first two digits (indicating the approximate row and column positions of the scale), and the third digit describing the number of sides. Individual comparisons were done by measuring the difference between each of the equivalent scale IDs. The main limitation with this procedure is the reliance on maintaining a consistent region in which to identify scales. Including or removing a scale from the compared identifiers will significantly change the probability of a true positive match. Additionally, since the scales do not follow a simple grid pattern, the row and column designations for scales further from the eye are more subject to errors. A similar method of analyzing scute patterns was used to identify alligators (Balaguera-Reina et al. 2017), but similarly neglected any topological information of the patterns. We propose performing individual identification for sea turtles by representing and comparing the scale patterns as mathematical graphs.

Mathematical graphs or networks have been used to describe many biological systems such as protein-protein interactions (Uetz et al. 2000), gene regulation (Arnone & Davidson, 1997), evolutionary history (Huson & Bryant, 2006), social interactions (Lusseau, 2003), and food webs (Briand & Cohen, 1984). As a result, pattern recognition on graphs is useful in many fields (Foggia et al. 2014). The performance of pattern recognition algorithms depends primarily on the choice of the algorithm used to compare the similarity of two given graphs. One of the most efficient classes of matching algorithms are spectral methods (Caelli & Kosinov, 2004b; Wilson & Zhu, 2008) which involve eigenvalue decomposition of the adjacency or Laplacian matrix of the graph and can be computed in polynomial time. A more flexible and versatile class of methods involves calculating the graph edit distance (Sanfeliu et al. 1983; Fischer et al. 2017), which is known to be NP-complete. Many polynomial algorithms have been suggested to approximate the graph edit distance, but only work for certain classes of graphs, or are not guaranteed to find global minima solutions (Justice & Hero, 2006). Finally, graph kernels (Gaüzère et al. 2012; Bai et al. 2015; Kondor & Pan, 2016) represent another general procedure in which the graph is converted to some vector which in turn is used as the input to a machine learning algorithm or support vector machine for further classification/matching.

The pattern of facial scales of sea turtles is specific to an individual and stable over its lifetime; thus, it can serve as the basis for automated facial recognition (Carpentier et al. 2016). We introduce an algorithm that accomplishes individual identification in two steps. First, an observer identifies scales (nodes), marks their positions, and collects the pattern of connections between them (edges). The nodes and edges define a map graph (Chen et al. 2002) for the face. We then generate a mapping between the nodes of each map graph using a coherent point drift algorithm (Myronenko & Song, 2010) on the scale positions, along with the eye and beak as points of reference. Second, we measure the similarity between the map graphs using a modified graph edit distance metric. Our algorithm correctly match turtles in a database 94% of the time. The main novelty and impact of this paper is to combine information on the positions and the pattern of connections of scales to perform individual identification.

## 2. Materials and Methods

We analyzed 164 photos of sea turtles, taken over a span of 10 years (2009–2019), in a range of locations and conditions (e.g. lighting, distance, angle), by different researchers, using different cameras. Included in this sample were green turtles, *C. mydas*, of both the black and green morphotype, and hawksbill turtles, *E. imbricata* (**Table 1**). All photos were of the right side of the face. Previously, these images had been judged to belong to at most 139 individuals, with 20 individuals photographed multiple times (20 viewed twice, 4 viewed 3 times, and 1 viewed 4 times). The database is maintained by J.P.M.P. and D.A.R. A map graph was generated from each image independently by K.K.R. and R.B.R.A. resulting in two sets of graphs for each image. The algorithms used to compare the images do not require any fitting, so no train/test/validation split is needed.

**Table 1:** List of previously identified number of individuals and species used in study

| **Species of Turtle** | **Number of Individuals** | **Number of Images** |
| --- | --- | --- |
| Green (Yellow Morphotype) | 31 | 39 |
| Green (Black Morphotype) | 101 | 113 |
| Hawksbill | 7 | 12 |
| Total | 139 | 164 |

### 2.1 Generating Map Graphs from Images

We used each image of a sea turtle face to construct an undirected graph, *G*= (*V, E*), where *V* is a set of nodes (or vertices) and *E* is a set of edges (or pairs of nodes). The nodes represent scales and the edges represent pairs of scales sharing a common border. We included two additional nodes: the eye and the tip of the beak, designated nodes 1 and 2, respectively. The eye is connected to its neighboring scales, but the beak is not connected to any other scales. We disregard the scales on the top of the head and the small scales on the neck and on the bottom of the turtle’s face. Thus, if we exclude node 2 (beak), graph *G* is the map graph (Chen et al., 2002) of the eye and scales. The resulting map graphs are unlabeled (except for nodes 1 and 2) and mostly planar, that is, they consist of edges that do not intersect each other. However, edges will intersect if three or more scales meet at one point (corner). The pixel coordinates of the nodes (at approximately the center of the scale) are also recorded as node attributes.

Some turtles have complex facial patterns that make scale identification difficult and neighborhood unclear, especially if the quality of the image is poor. In addition, light conditions may create artifacts that can be mistaken for scale borders. These problems make graph generation difficult to automate. As a result, one of the main sources of error in our approach is inconsistent generation of graphs. An example is shown in **Figure 1** with additional sample images of the Hawksbill and Yellow morphotype are given in **Figure S1**.

**Figure 1.**
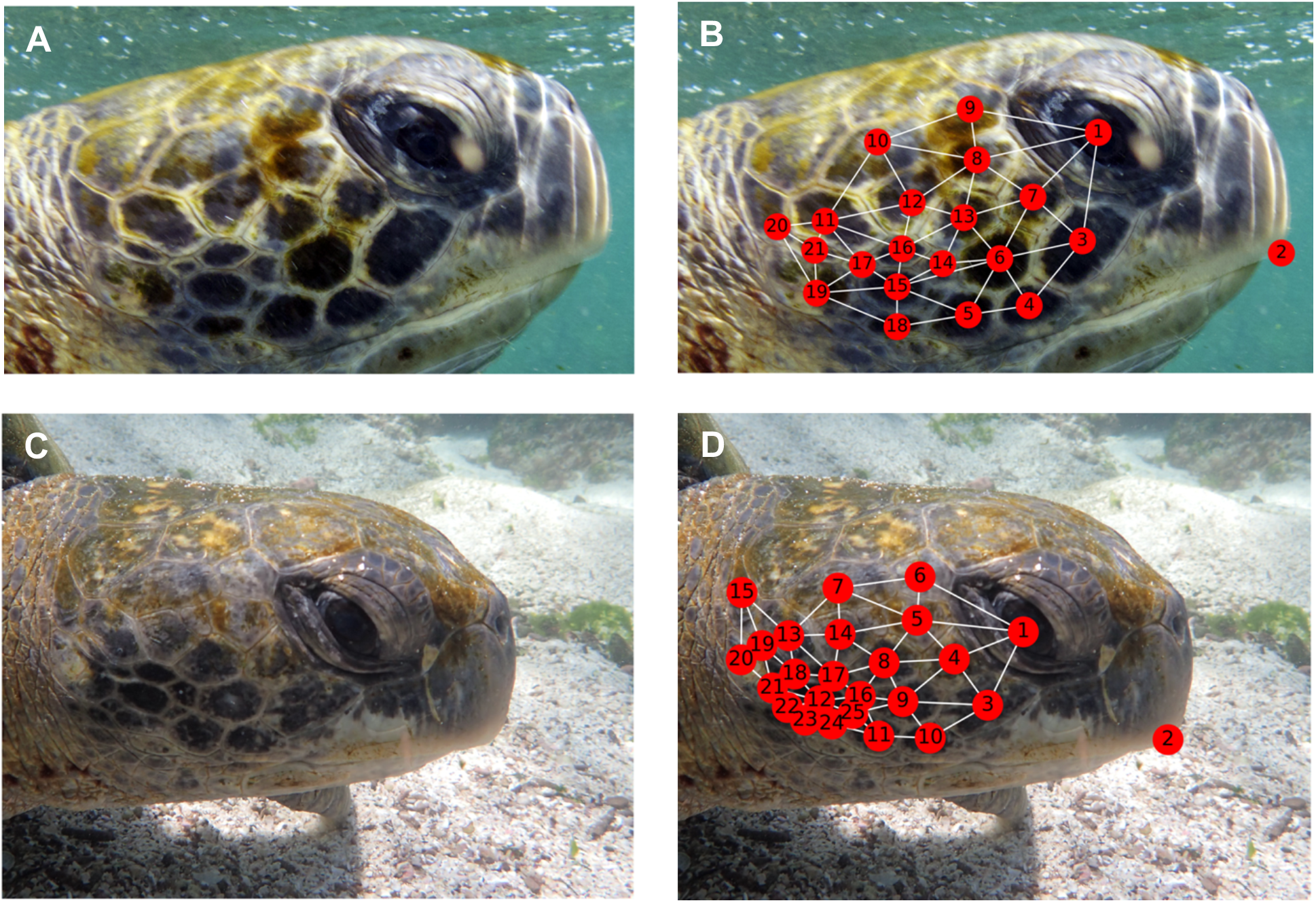
Errors in reconstructing face graphs. (A) and (C) are two photographs of the same turtle taken under different conditions and analyzed by the same researcher at different times. (B) and (D) are the associated map graphs generated for each image. Node 1 is the eye and node 2 is the tip of the beak. The remaining nodes represent scales; their numbers indicate the (arbitrary) order in which they were selected. Edges indicate whether scales and/or the eye are neighbors. See Figure 3 for detailed comparisons of the map graphs in (B) and (D).

To evaluate the effect of these variations in graph reconstruction, two sets of the turtle graphs were obtained by two researchers independently generating one graph from each image. When multiple images were thought to belong to the same turtle (based on earlier evaluations), each investigator made sure that their graphs were generated independently, at least two hours apart, to minimize the impact of memory on graph generation.

### 2.2 Graph Topology Metrics

The graph topology of the scale structure guarantees three invariances. First, invariance to translation; the picture of the turtle may occur anywhere on the image. Second, invariance to rotation of the turtle’s face in three dimensions. Third, invariance to lighting and visibility conditions. The first two properties show that the graph is a global property of the turtle’s face. Ultimately, the graph is insensitive to how the photo is taken, provided the relevant scales and borders are clearly visible. For a given graph *G*, there are many global measures of the size and connectivity. We consider 11 different metrics of the size and connectivity of the graphs (see **Table S1**) and perform principal component analysis (PCA) on these metrics to evaluate the variation in graph topology within and among species.

### 2.3 Graph Spectrum

Two metrics we used to compare the topology of two graphs were based on the spectrum of their adjacency and Laplacian matrices. The adjacency matrix (**A**) of graph *G* is a square matrix whose element *A*_*ij*_is 1 if nodes *i*and *j*are connected and 0 otherwise (*A*_*ii*_ = 0 ∀_*i*_). The Laplacian matrix (**L**) of *G* is defined as

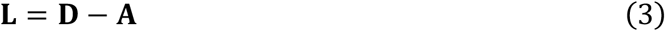

where Dis the degree matrix of *G*, a diagonal matrix, whose element *D*_*ii*_ is the degree of node *i*, that is, the number of edges connecting to that node. The spectrum of the **A** or **L** matrices is the set of eigenvalues *λ*_*i*_. We calculate the following distance (*S*) between the spectra of the two graphs:

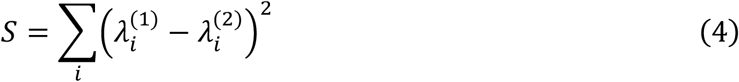

where 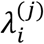 is the *i*^th^ eigenvalue of graph *j* and *i* indexes over the minimum number of eigenvalues needed to capture 90% of the total power of the spectrum, typically *min*(*n*_2_, *n*_2_) − 1 where *n*_*j*_ is the number of nodes in graph *j*. Two isomorphic graphs will have *S* = 0, but *S* = 0 does not guarantee isomorphism. *S* does not make any assumptions about which nodes (i.e. scales) match between two turtle faces.

### 2.4 Coherent Point Drift with Scale Positions

The nodes were selected to lie approximately on a plane, so the positions of the nodes in all possible orientations or perspectives are related by an affine transformation. Solving for the affine transformation that aligns a series of unmarked points has been thoroughly studied in computer science as the point set registration problem (Besl & McKay, 1992). Considering the two sets of unlabeled points (nodes) *G* and *H* from two turtle faces with points *G*_*n*_ and *H*_*m*_, respectively, related by some non-rigid, affine transformation *T*:

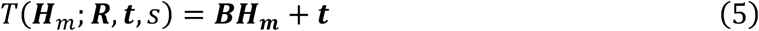

where ***B*** is an affine transformation matrix (including rotation ***R*** and scale, *s*), ***t*** is a translation vector, and *H*_*m*_ refers to point *m* in graph *H*. The coherent point drift (CPD) algorithm proposed by Myronenko & Song, 2010 is applied to both solve for the ideal transformation and determine a mapping between two sets of nodes. As defined in the CPD algorithm, the probability of correspondence between nodes *G*_*n*_ and *H*_*m*_ can be calculated as

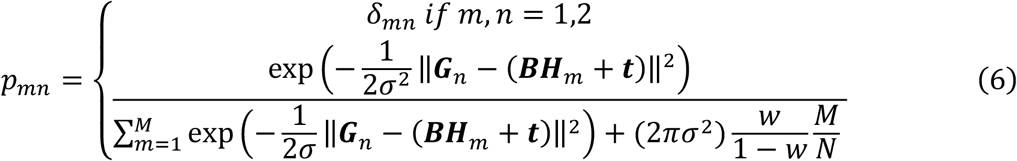

where *δ*_*mn*_is the Dirac Delta function, *w* is a weight parameter (we assume *w* = 2), and *M* and *N* are the total numbers of points in the *G* and *H* graphs, respectively. Points 1 and 2 are always identified as the eye and tip of the beak in every graph, so we set the probability of correspondence of those two points to their corresponding points in the other graph to 1 and the probability of correspondence of those two points to all other points to 0. By fixing these probabilities, we bias the estimation to align the eye and beak of the turtle at the expense of finding a transformation that aligns more nodes. We believe this to be an acceptable trade-off because the physical location of the scales is important in the identification of the turtle and prevents the algorithm from finding a more accurate matching in which the whole point cloud is arbitrarily rotated or translated. An iterative approach is then used to calculate a transform *T* that maximizes velocity coherence, which is a type of smoothness on the transformation (Yuille & Grzywacz, 1988). After applying the CPD algorithm, the most likely mapping between nodes is calculated from the maximum values of *p* in each row.

Based on the mapping calculated using the CPD algorithm, we can additionally estimate the covariance between two point sets *G* and *H* as:

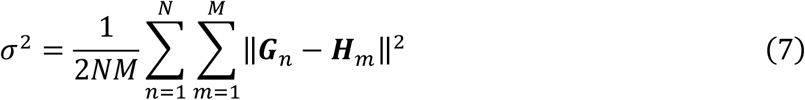

which is a measure of distance between the two sets of coordinates.

### 2.5 Graph Edit Distance

The graph edit distance (GED) is a measure of the minimum number of modifications that must be made to convert one graph into another by adding or subtracting nodes and edges (substitution operations). Here we compare two quadratic-time approximations of GED: Bipartite Matching (Fischer et al., 2017; Riesen & Bunke, 2009) and Greedy Edit Distance (Fischer et al. 2015). These two approximations to GED do not make any assumptions about which nodes (i.e. scales) match between two turtle faces.

We also introduce a modified position-corrected graph edit (PGED) needed to make *G* isomorphic to *H*:

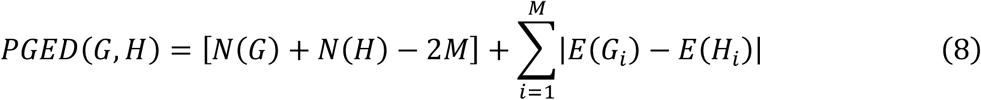

where *N*(*G*) is the number of nodes in a graph, and *E*(*G*_i_) is the number of edges (i.e., the degree) of node *G*_i_ of graph *G. M* is the number of nodes that were matched between *G* and *H* by the CPD algorithm (Section 3.4). Thus, *PGED*(*G, H*) is not necessarily equal to *PGED*(*H, G*) which means PGED is not strictly a distance metric but is an approximation of the GED which is a true distance (Bougleux et al., 2017). Equation 8 is a count of the number of nodes that have no equivalent in the other turtle (the expression in square brackets) plus the absolute differences in the degrees of matched nodes. Using PGED as the distance metric in the ranking algorithm penalizes graphs with larger differences in number of nodes and where the matched nodes have different degrees. Note, however, that the comparison of the degrees of two nodes ignores which nodes are actually connected to each other.

## 3. Results and Discussion

### 3.1 Matching Via Isomorphism

For a given graph of a turtle *G*, and a database of other turtle graphs *ℋ* we wish to define a distance function between *G* and each graph *H* in *ℋ, D*(*G, H*), that is minimized if graphs *G* and *H* are from the same turtle (ideally, equal to 0), and greater than 0 otherwise. The ‘simplest’ form of *D* would be an isomorphism test between *G* and *H*: if the two graphs are isomorphic, they belong to the same turtle, and if they are not isomorphic, they belong to different turtles. Even ignoring the computational complexity of such an algorithm (testing for graph isomorphism is an NP-complete problem), such an approach would result in many false negatives because it assumes differences in graphs are only a result of actual differences between individuals and not errors in graph generation. Although the graph structure itself is a robust description of the scale structure, there is a large source of variation associated with uncertainty in determining the graph from different images, as well from differences of opinion by different observers or even the same observer at different times.

An example is shown in **Figures 1B and 1D** where two images of the same turtle, taken at different times, and annotated by the same person, result in different facial map graphs. We know *G* and *H* belong to the same turtle, but they could differ in the number of nodes, and possible connection of edges, and therefore need a more general distance metric than the Boolean isomorphism to compare *G* and *H*. The average difference in number of nodes between the two sets of the same images (generated by different researchers) was 1 ± 2.2 and the average difference in total number of edges was 2.6 ± 5.5 (mean ± two standard deviations) based on the 164 turtles viewed by both researchers. For a given turtle image, the two researchers produced graphs with identical numbers of nodes and edges only 22% of the time. Thus, an effective metric must be able to tolerate errors in graph generation involving on the order of 1 node and 3 edge differences.

### 4.2 General Graph Topology

We consider the distribution of 11 properties of the turtle graphs and perform principal component analysis (PCA) to determine if the individuals and species are separable by a linear combination of these properties (see Supplemental Information **Table S1** for the full list of properties). The first two principal components capture 63% and 13% of the variance, respectively, with loadings shown in **Figure S2. Figure 2A** shows that hawksbill and green sea turtles are well discriminated along PC1. This indicates that the two species have distinct characteristic facial map graph topologies. The yellow and black morphotypes of green turtles, however, do not form distinct clusters based on PC1 and PC2 (**Figure 2A**).

**Figure 2.**
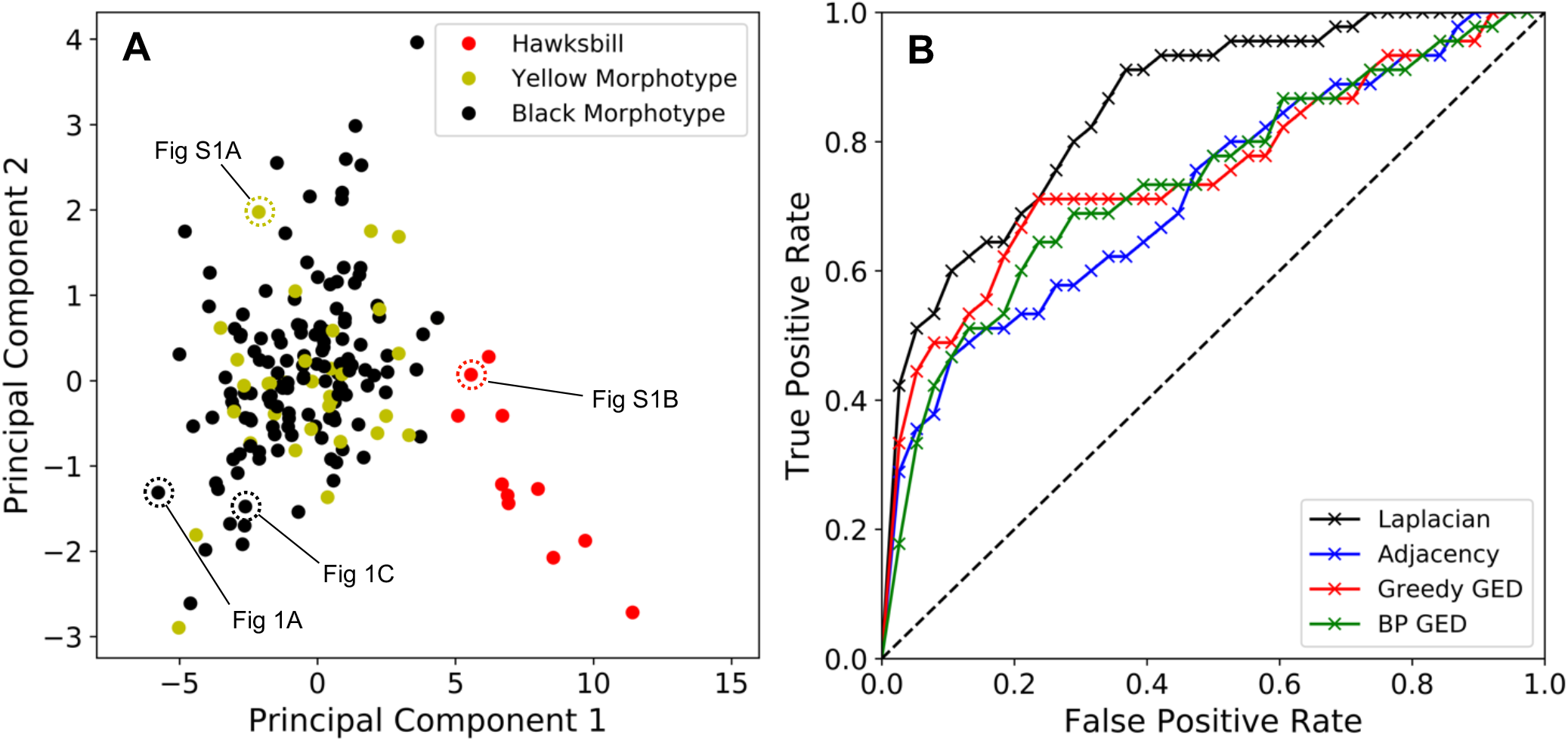
The topology of the face graph can be used to distinguish two species of sea turtle. (A) Each turtle face graph projected along first two principal components of graph properties. Colors indicate species. Hawksbills are clearly separated from the yellow and black morphotypes along PC1. Points corresponding to the black morphotype turtles in **Figures 1A and 1B** and for the yellow morphotype and hawksbill shown in **Figures S1A** and **S1B** are annotated. (B) ROC curves using the spectrum similarity and GED methods to compare turtle graphs. A false positive rate of 0.12 corresponds to only considering the top 20 candidate matches.

Although PCA may be adequate for species identification, it is insufficient for the purposes of individual identification because even different turtles have similar map graphs, as the distance between individuals in this latent space is greater than the average distance between any two points. For the two images of the turtles shown in **Figure 1A/C** as an example, their corresponding graphs are not close to each other along either principal component, so any form of nearest neighbor classification, or clustering, would result in many false positive identifications. Since these principal components only capture general properties of graph topology, additional information regarding the position and local environments of nodes are necessary to distinguish individuals.

### 4.3 Matching Via the Graph Spectrum and Graph Edit Distance

Another measure for the general topology of a graph is the spectrum and includes information on the algebraic connectivity described in section 3.3. Using the distance metric defined in equation 4, we compute all pairwise similarities within each of the two sets of image graphs generated by the two researchers (26,732 comparisons for each set). We then calculated the rank of all known duplicate turtle comparisons, i.e. rank between two different images of the same turtle. If we consider a series of cutoff ranks *r*_*c*_= 0,1, … 163, the false positive rate is the cutoff normalized to the total number of comparisons per image, i.e. 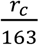, which is in the interval [0,1]. The true positive rate is the normalized number of matches with rankings less than the cutoff rank. In Figure 2B we plot the true positive rate as a function of false positive rate, i.e., the receiver operating characteristics (ROC) curve, and the corresponding area under the curve (AUC) is a measure of the overall performance of an algorithm with AUCs closer to 1 indicating better algorithm performance.

Using the Laplacian spectrum, the average rank between two different photos of identical individuals is 27.0 with only 60% of true known positive matches captured within the top 20 candidate images (false positive rate = 0.12), i.e., if a user were manually searching the top 20 most similar images, there is a 60% chance of the duplicate individual image appearing if such a match exists in the database (**Figure 2b, Table 2**). A similar proportion was observed when analyzing the second set of independently generated images with the average rank of known positive matches of 21.5 and 66.7% of true matches caught within the top 20. Using the adjacency spectrum resulted in a worse performance with the average rank of known positive matches at 51.9, and 46% of true matches occurring in the top 20 candidates. The ROC curves for the Laplacian and adjacency spectra are shown in **Figure 2b**, each with an AUC of 0.85 and 0.72, respectively. Both the Greedy and Bipartite algorithms for estimating the GED between two graphs perform worse than the Laplacian eigen spectrum with average ranks of known positive matches at 45.5 and 49.2, respectively and only 49% and 47% of true matches identified in the top 20 (**Table 2**). The poor performance of these GED methods is also reflected in the lower AUC values of 0.75 and 0.74 for Greedy and Bipartite respectively, which are of the same order as the adjacency spectrum method (**Figure 4, Table 2**).

**Table 2:** Performance of different graph comparison metrics compared by AUC (area under the ROC curve), and fraction of true positive matches found within the top 20 proposed matches.

| **Metric** | **AUC** | **Average Match Rank** | **% Caught < 20** |
| --- | --- | --- | --- |
| Laplace Eigen Similarity | 0.85 | 27.00 | 60.0 |
| Adjacency Eigen Similarity | 0.72 | 51.93 | 46.7 |
| Bipartite GED | 0.74 | 49.22 | 46.7 |
| Greedy GED | 0.75 | 45.47 | 48.9 |
| Point Covariance | 0.87 | 23.31 | 71.1 |
| Delta Nodes | 0.92 | 14.11 | 82.2 |
| Delta Edges | 0.92 | 13.42 | 86.7 |
| Position GED | 0.94 | 9.20 | 91.1 |
| Position GED + Covariance | 0.90 | 16.31 | 82.2 |

The relatively poor performance of the spectrum and GED measures can be attributed to the homogeneity of the size and topology of the graphs among turtles—the standard deviations of the numbers of nodes and edges are 1.9 and 4.2, and the graphs are mostly planar being composed of a series of non-overlapping triangles. Thus, completely different groups of nodes in different turtles can show very similar patterns of connectivity.

### 4.4 Scale Positions

Here we investigate if the relative positions of the scales, independent of their map graph, are sufficient to identify an individual. Even if *G* and *H* are not isomorphic, there exists some mapping between a subset of nodes in *G* and *H* that match the same scales in each graph which may be calculated entirely based on the coordinates of the nodes (and ignoring the connectivity information). Nodes 1 and 2 will always be fixed markers, but the numbering of the nodes is arbitrary after nodes 1 and 2. We use the coherent point drift (CPD) algorithm described in section 3.3 to find the correspondence between nodes in *G* and nodes in *H* based on the coordinates of the nodes while maintaining the overall topology for the graph (**Figure 3A/B**). The CPD algorithm assigns a probability that node *n* from *G* is homologous to node *m* from *H*.

**Figure 3.**
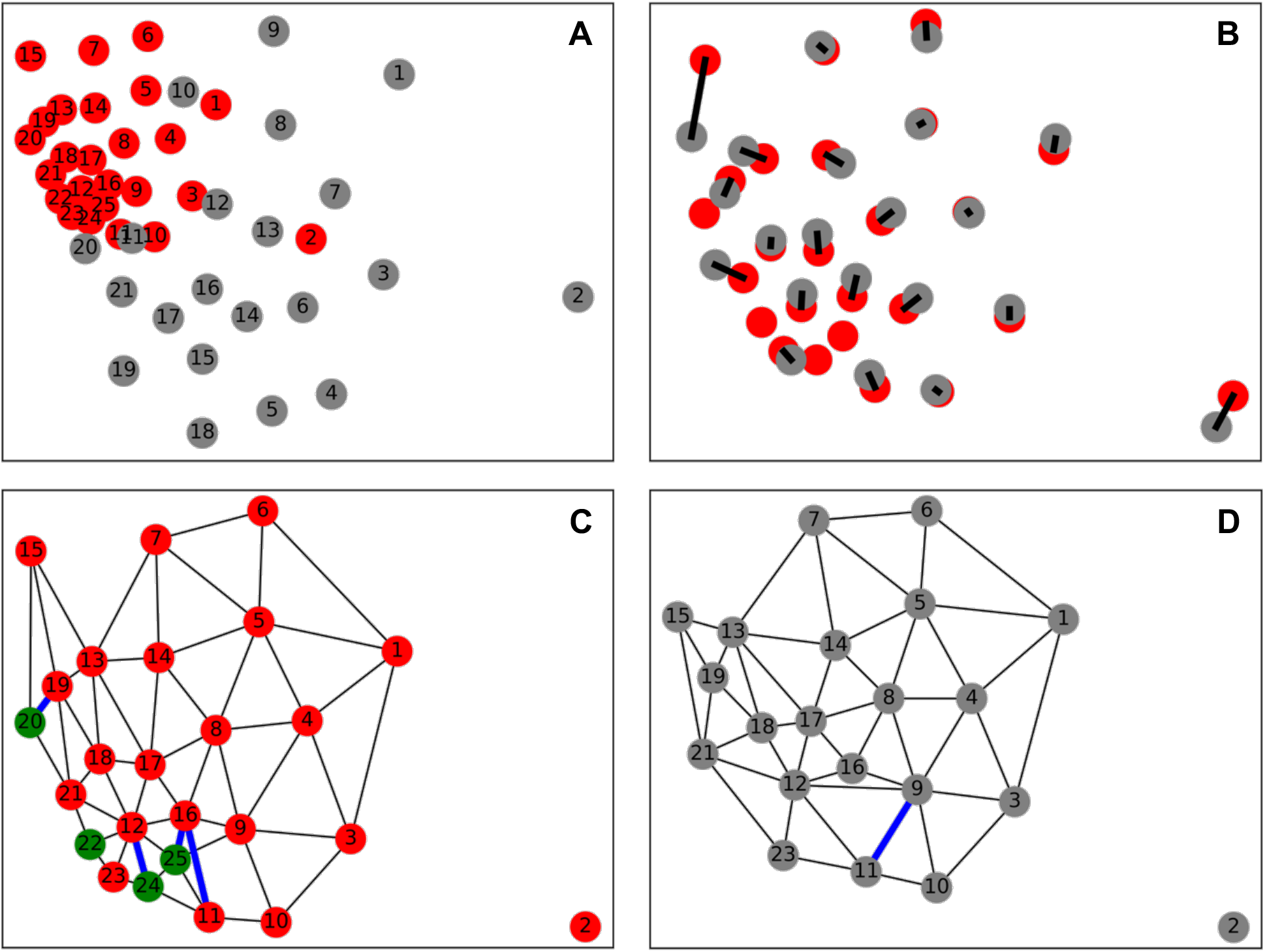
Outline of the various steps needed to compare two face graphs using the position-corrected graph edit distance. (A) Original coordinates of the nodes in the face graphs. Numbers indicate the order in which the node was recorded. Nodes from Figure 1A/B are in grey, nodes from Figure 1C/D are in red. (B) Calculated transformation by the coherent point drift algorithm (numbers omitted for clarity), with mapping between nodes calculated as equivalent connected by a line. (C) and (D) show the two graphs after point registration. Nodes with no mapping are highlighted in green (4 total) and edge differences of the remaining matched nodes highlighted in blue (5 total) which sum to determine the *GED* between two graphs (*PGED* = 9). The nodes in (D) are renumbered to match the nodes in (C) based on the calculated mapping.

**Figure 4.**
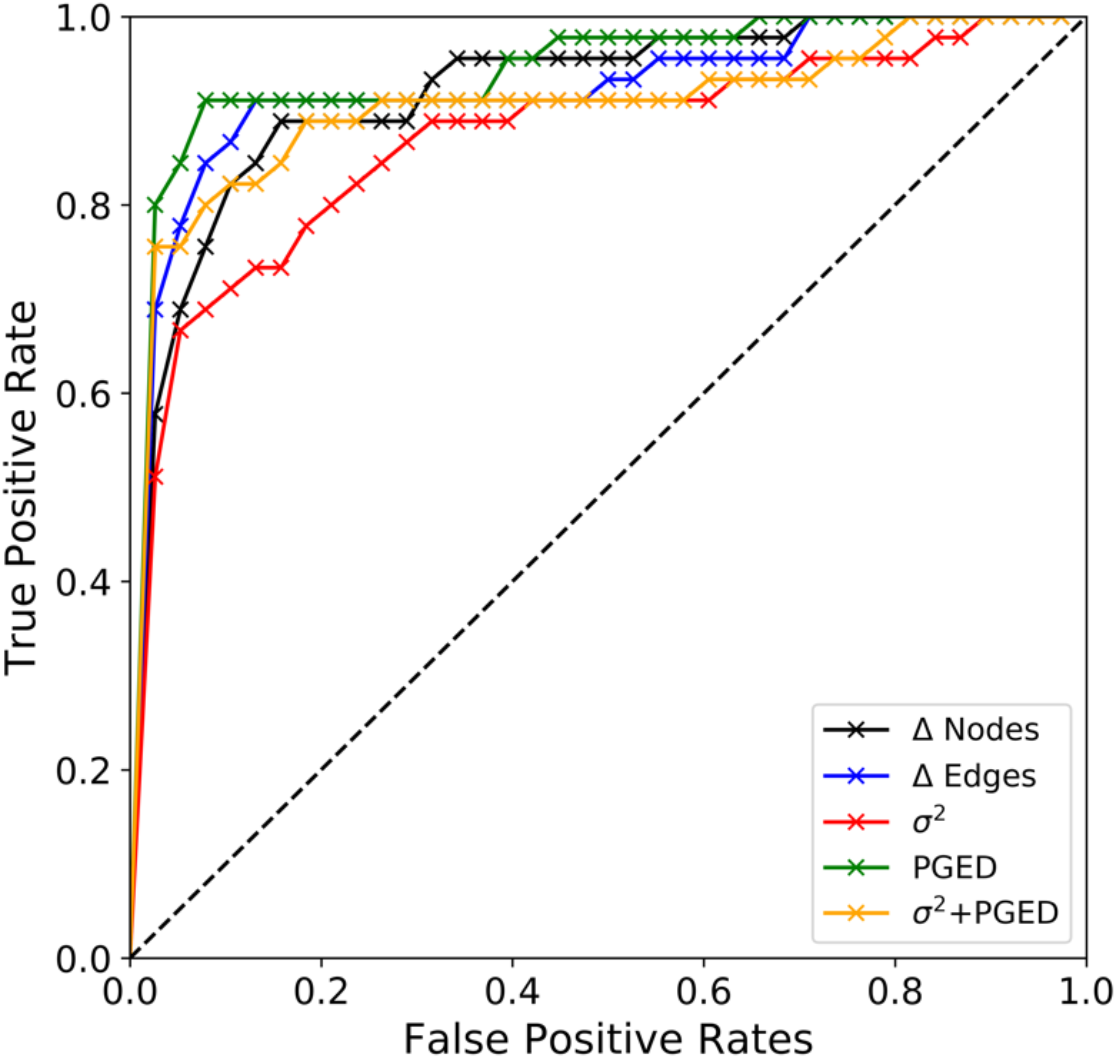
ROC characteristics using different metrics for comparing similarity between two turtle graphs. The new PGED algorithm performs the best with an AUC of 0.94.

We note that the transform found in the CPD algorithm (and therefore the covariance, σ^2^) is not commutative with respect to the choice of the reference graph, i.e. σ^2^(*G, H*) ≠ σ^2^(*H, G*). We set the graph *G* we are searching for as the reference graph to maintain consistency (i.e., calculate the transformation for a graph in the database to match a new graph). This approach results in a normal distribution for σ^2^. Changing the reference with each comparison and using the database graph as the reference significantly changes the distribution of σ^2^ increasing the fraction with low covariances (**Figure S3**). Ranking turtles based on the covariance, after calculating the most likely mapping between the two sets of node positions, improves the accuracy of the matching algorithm with the average rank of a true positive match decreasing to 23.3 and 71% of true positives captured in the first 20 comparisons (**Table 2**). The resulting ROC curve using only the covariance is shown in **Figure 4** with an AUC of 0.87.

The CPD algorithm will provide false positive results when the two graphs have significantly different numbers of nodes. If the numbers of nodes in *G* and *H* are not of similar magnitude, it becomes much easier to fit the point distribution of *H* to *G* and vice versa. Using only the covariance of equation 7 there is no penalty associated with the number of nodes that are discarded and do not have an equivalent node in the other graph. Additionally, the information about the shape of the node is lost in the covariance. For example, the eye is always a mappable point by definition, but the number of neighboring scales is not considered.

### 4.5 Position-Corrected Graph Edit Distance

We introduce a new approach combining the GED and CPD algorithms. After determining the optimal mapping of nodes based on the CPD algorithm, we calculate the GED between the graphs using equation 8 (**Figure 3C and 3D**). This position-corrected GED (PGED) approach produced the best results with the lowest average rank of known true positive results at 9.2 and 91% possible true positives captures within the first 20 images.

Using either just the differences in nodes mapped, or the edges for the mapped nodes did not perform as well with average rank of true positives at 14.1 and 13.4 respectively. While the difference in nodes measures how many identifiable scales there were, the difference in number of edges measures an uncorrelated effect of the shape of the scale, which is why the sum of two provides a better performance than either alone. Using the sum of the covariance and the PGED did not improve the accuracy of the model because the spread in covariance values was much larger than the uncertainty in the differences in edges and lowered the algorithm’s performance. The ROC curves for each of these metrics are shown in **Figure 4**. For the second curated dataset, the PGED was similarly the best performing metric with the average rank of duplicates at 12.9 and 84% presented in the top 20 individuals and AUC of 0.94 (**Table 2**). The PGED metric performed less well on the second dataset, but still outperformed the other metrics.

While the vast majority of the true positive ranks were less than 10, there appeared to be 6 outlier comparisons where the known true positive ranked near the bottom with ranks of 81, 121, 163, 120, 72, and 73 respectively. When we looked at the calculated highest ranking matches for each of these cases, we found that the top identified candidate in each of the first four cases was actually another true positive that had been erroneously considered as a different individual based on previous analysis. In the second dataset, there were a similar 8 outliers with true positive ranks of 46, 58, 134, 45, 120, 44, 119, and 116, which encompass the same 6 previously mentioned individuals that contained unknown positive matches. However, the rank of these unknown positive matches in the second dataset was slightly lower with an average rank of three compared to one in the first dataset. Reevaluating all 163 images (only considering top 5 matches), we found an additional 20 duplicate individuals that were missed by previous analyses (in addition to the 4 mentioned above). Therefore, the number of individuals had been overestimated by at least 14.7% in the database.

## 5 Conclusion

We introduced an algorithm for individual identification of sea turtles using a map graph representation of the facial scale structure. To account for variations in orientation and image quality, we used a modified coherent point drift (CPD) algorithm to generate a mapping between nodes of two graphs. We then calculated a modified graph edit distance between mapped nodes as a distance metric between two graphs. Ranking based on only the position-corrected graph edit distance provided the lowest average rank (9.2) than if the covariance between graphs is included or with any of the other distance metrics we considered (**Table 2**). Additionally, our method was able to identify at least 24 unknown matches between turtles in our database that had been mislabeled as different individuals by experienced observers.

The main limitation of our approach is the manual process of generating the graphs from each image. We estimate the time needed to annotate the graph is approximately 30 seconds and is negligible compared to the time needed to acquire the photo of the turtle in the first place, which requires swimming, searching, and diving. The high variation in the colors of scales, lighting conditions, and visibility makes automation of this process difficult, but might be approached using image segmentation machine learning models (Papandreou et al. 2015).

We believe our approach will provide a robust foundation for the use of facial recognition for individual identification of turtles and can be used by scientists all around the world to collect and share information related to sea turtle populations. This information can also be collected with citizen science efforts where naturalist guides, local students, fishermen, and tourists can collect and share this critical information in vast areas like the Galápagos Marine Reserve and other marine reserves.

This matching algorithm may also assist in individual identification for other species whose scales or spot patterns are unique such as whale sharks (Brooks et al. 2010), giraffes (Halloran et al. 2015), and leopards (Miththpala et al. 1989) where connectivity and topology information is often discarded in favor of just the positions of key features. More research is needed into machine learning approaches to wildlife population surveys.

## Acknowledgements/Ethics

The authors declare no conflicts of interest.

Data collection for this project was authorized by the Galápagos National Park (GNP) Service (Permit # PC-21-19 to DAR and JPMP) and approved by UNC-Chapel Hill & Universidad San Francisco de Quito (USFQ) Galápagos Science Center (GSC) ethics and animal handling protocols.

KR and LG acknowledge the NASA Space Technology Research Fellowship (grant # 80NSSC17K0148); RA acknowledges the National Science Foundation (grant # DEB-2014566). The University of Houston (UH) College of Natural Sciences and Mathematics, the Department of Biology and Biochemistry, and the Honors College provided financial support. We are grateful to the GNP authority for permitting data collection for this project; to the UNC-USFQ GSC for facilitating fieldwork and lab space; and to the volunteers who assisted during data collection in Galápagos. We thank Lydia Golightly for help with preliminary data collection in Houston. Special thanks to the ranger Jason Castañeda, the GNP technical office at San Cristóbal Director Maryuri Yépez, and the GalápaGo! UH team Tony Frankino, Marc Hanke, Ann Cheek, and Rebecca Zufall.

## Author Contributions

KR and RA conceived the ideas and designed methodology; DAR and JPMP collected the data; KR and RA and LCG analyzed the data; KR and RA led the writing of the manuscript. All authors contributed critically to the drafts and gave final approval for publication.

## Code and Data Availability

The code and data will be made available through GitHub at the time of publication.

## Supplemental Information for

### Automation Detection for Side of Face

Although in this study we only considered the right side of the face, there’s no reason individual identification could not be done on the left side. It is important for the matching algorithm to only match graphs generated from the same side of the face, so here we present a simple method for determining which side of the face the image was taken using the same graph. Automating this step increases the overall efficiency of matching and removes one possible source of human error. From the turtle graph we label the positions of the eye and beak as *X*_1_ = (*x*_1_, *y*_2_) and *X*_2_ = (*x*_2_, *y*_2_), respectively, and a third point as the average coordinate of the other nodes:

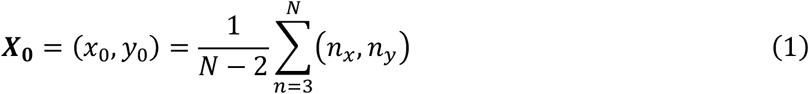

where *N* is the number of nodes. We then can define two vectors, *V*_1_ = (*x*_1_ − *x*_0_)*î* + (*y*_1_ − *y*_0_)*ĵ* and *V*_2_ = (*x*_2_ − *x*_1_)*î* + (*y*_2_ − *y*_1_)*ĵ* which point from the center of the scales to the eye, and from the eye to the beak respectively. The cross product of the two vectors has only a k component, whose sign is determined by the side of the face:

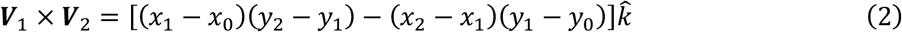

The right side of the face would result in a positive sign for equation 2, and the left side of the face would have a negative sign. These signs correspond to a clockwise and counterclockwise rotation between the vectors respectively, and can be easily verified in **Fig. 1** with the *right-hand-rule*. This inequality is invariant under any rotation of the image.

**Fig. S1.**
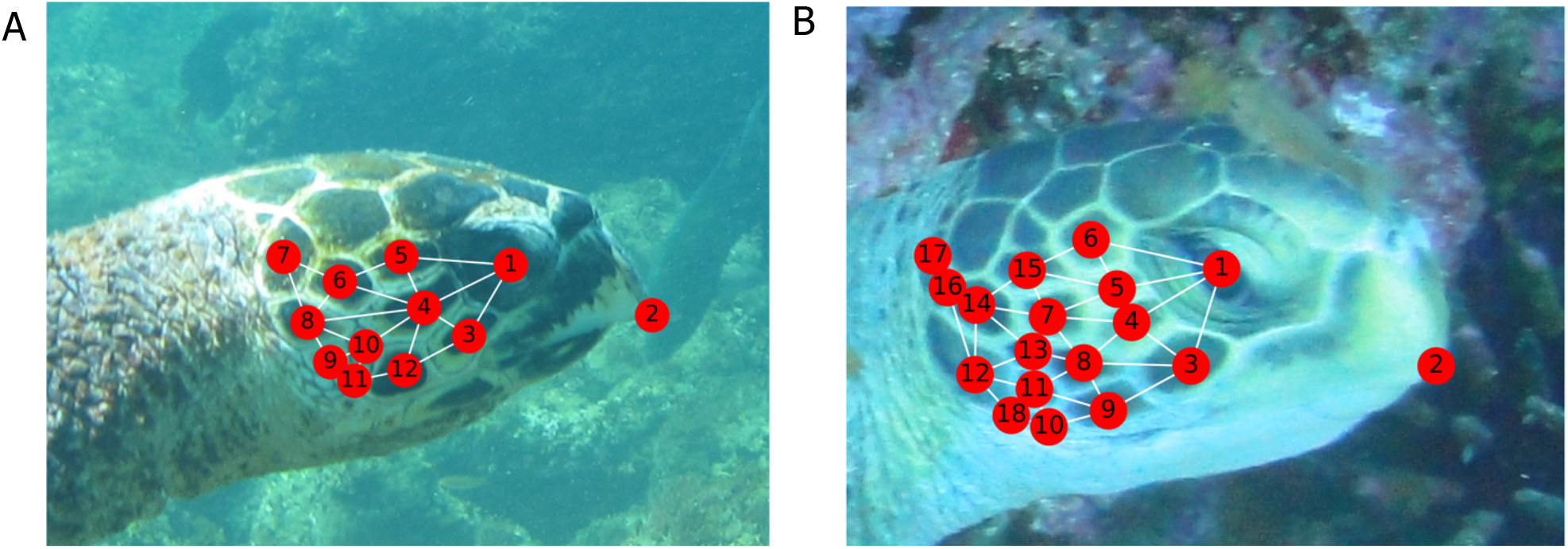
Image and associated graph of a (A) hawksbill species and (B) a green sea turtle yellow morphotype.

**Fig. S2.**
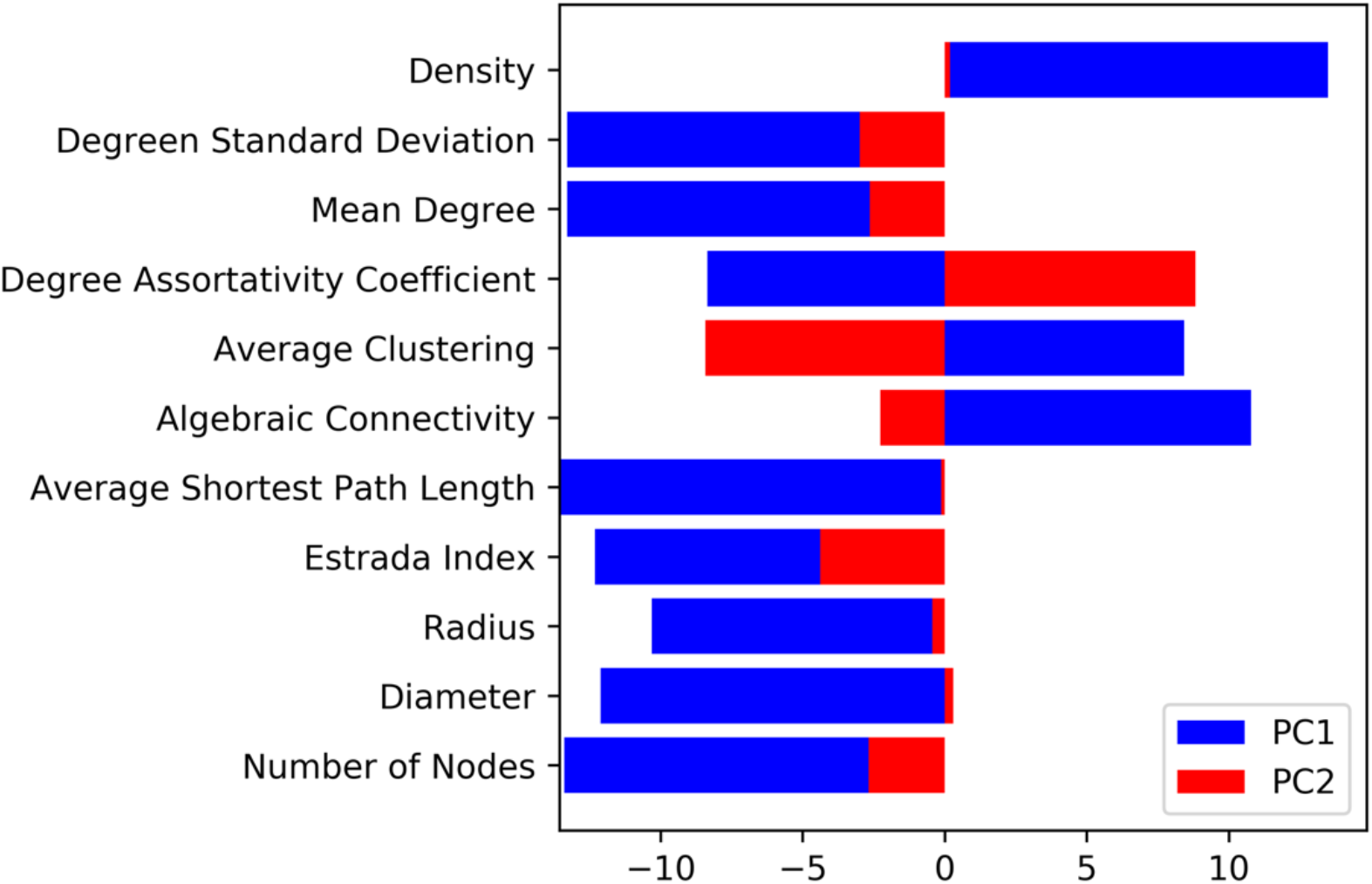
Loadings of the first two principal components.

**Fig. S3.**
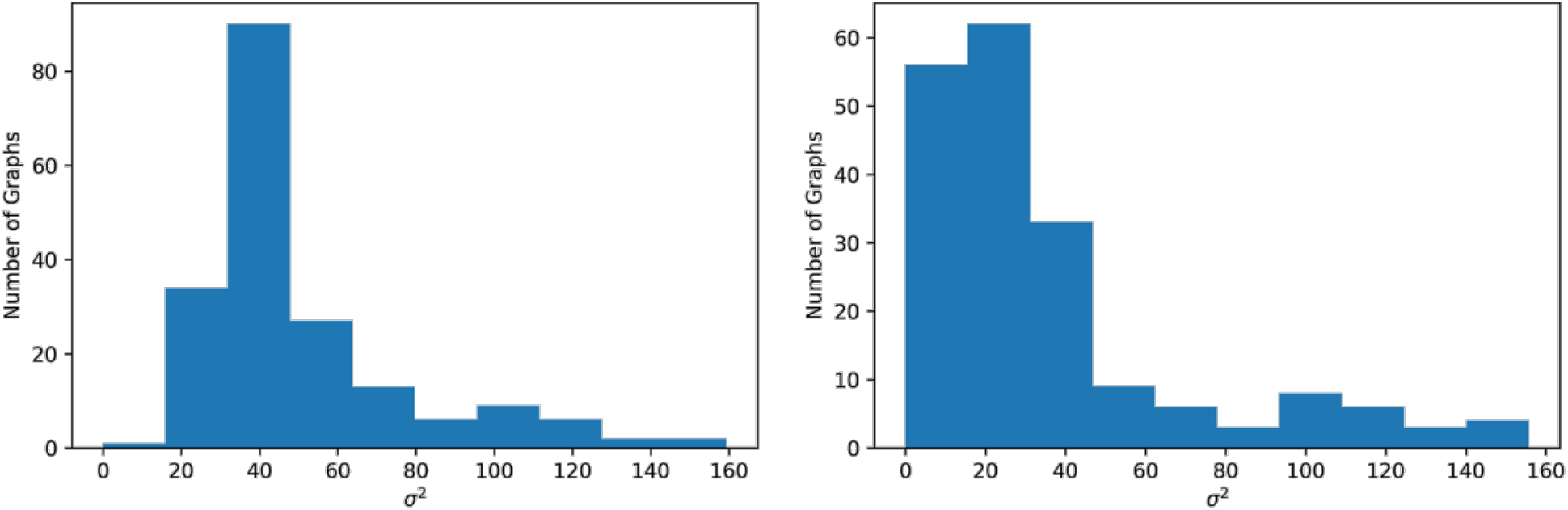
Histogram of covariances between a turtle graph *G* and other graphs *H*_*i*_ using (A) *G* as the reference graph, and (B) *H*_*i*_as the reference graph.

**Table S1:** List of Graph Properties considered in PCA analysis

| Property | Include Node 2? (Beak) |
| --- | --- |
| Number of Nodes | Yes |
| Graph Diameter | No |
| Graph Radius | No |
| Estrada Index | No |
| Average Shortest Path Length | No |
| Algebraic Connectivity | No |
| Average Clustering | Yes |
| Degree Assortativity Coefficient | Yes |
| Mean Node Degree | Yes |
| Density of Graph | Yes |
| Standard deviation of node degrees | Yes |

## References

Arnone, M. I., & Davidson, E. H. (1997). The hardwiring of development: Organization and function of genomic regulatory systems. Development, 124(10), 1851–1864.

Arzoumanian, Z., Holmberg, J., & Norman, B. (2005). An astronomical pattern-matching algorithm for computer-aided identification of whale sharks Rhincodon typus. Journal of Applied Ecology, 42(6), 999–1011. https://doi.org/10.1111/j.1365-2664.2005.01117.x

Bai, L., Rossi, L., Torsello, A., & Hancock, E. R. (2015). A quantum Jensen-Shannon graph kernel for unattributed graphs. Pattern Recognition. https://doi.org/10.1016/j.patcog.2014.03.028

Balaguera-Reina, S. A., Venegas-Anaya, M., Rivera-Rivera, B., & III, L. D. D. (2017). Scute Patterns as an Individual Identification Tool in an American Crocodile (Crocodylus acutus) Population on Coiba Island, Panama. Journal of Herpetology, 51(4), 523–531. https://doi.org/10.1670/17-023

Besl, P. J., & McKay, N. D. (1992). A Method for Registration of 3-D Shapes. IEEE Transactions on Pattern Analysis and Machine Intelligence, 14(2), 239–256. https://doi.org/10.1109/34.121791

Beugeling, T., & Branzan-Albu, A. (2014). Computer vision-based identification of individual turtles using characteristic patterns of their plastrons. Proceedings - Conference on Computer and Robot Vision, CRV 2014, 203–210. https://doi.org/10.1109/CRV.2014.35

Bougleux, S., Brun, L., Carletti, V., Foggia, P., Gaüzère, B., & Vento, M. (2017). Graph edit distance as a quadratic assignment problem. Pattern Recognition Letters, 87, 38–46. https://doi.org/10.1016/j.patrec.2016.10.001

Briand, F., & Cohen, J. E. (1984). Community food webs have scale-invariant structure. Nature, 307(5948), 264–267. https://doi.org/10.1038/307264a0

Brooks, K., Rowat, D., Pierce, S. J., Jouannet, D., Vely, M., & Al, K. B. E. T. (2010). Seeing spots: photo-identification as a regional tool for whale shark identification, 9(2), 185–194.

Caelli, T., & Kosinov, S. (2004a). An eigenspace projection clustering method for inexact graph matching. IEEE Transactions on Pattern Analysis and Machine Intelligence. https://doi.org/10.1109/TPAMI.2004.1265866

Caelli, T., & Kosinov, S. (2004b). Inexact Graph Matching Using Eigen-Subspace Projection Clustering. International Journal of Pattern Recognition and Artificial Intelligence, 18(3), 329–354. https://doi.org/10.1142/S0218001404003186

Calmanovici, B., Waayers, D., Reisser, J., Clifton, J., & Proietti, M. (2018). I 3 S Pattern as a mark recapture tool to identify captured and free-swimming sea turtles: An assessment. Marine Ecology Progress Series, 589(February), 263–268. https://doi.org/10.3354/meps12483

Carpentier, A. S., Jean, C., Barret, M., Chassagneux, A., & Ciccione, S. (2016). Stability of facial scale patterns on green sea turtles Chelonia mydas over time: A validation for the use of a photo-identification method. Journal of Experimental Marine Biology and Ecology, 476, 15–21. https://doi.org/10.1016/j.jembe.2015.12.003

Chen, Z.-Z., Grigni, M., & Papadimitriou, C. H. (2002). Map Graphs. J. ACM, 49(2), 127–138. https://doi.org/10.1145/506147.506148

Conte, D., Foggia, P., Sansone, C., & Vento, M. (2004). Thirty years of graph matching in pattern recognition. International Journal of Pattern Recognition and Artificial Intelligence, 18(3), 265–298. https://doi.org/10.1142/S0218001404003228 den Hartog, J., & Reijns, R. (2019). I3S Pattern (4.0.2).

Duyck, J., Finn, C., Hutcheon, A., Vera, P., Salas, J., & Ravela, S. (2015). Sloop: A pattern retrieval engine for individual animal identification. Pattern Recognition, 48(4), 1059–1073. https://doi.org/10.1016/j.patcog.2014.07.017

Fischer, A., Riesen, K., & Bunke, H. (2017). Improved quadratic time approximation of graph edit distance by combining Hausdorff matching and greedy assignment. Pattern Recognition Letters, 87, 55–62. https://doi.org/10.1016/j.patrec.2016.06.014

Fischer, A., Suen, C. Y., Frinken, V., Riesen, K., & Bunke, H. (2015). Approximation of graph edit distance based on Hausdorff matching. Pattern Recognition. https://doi.org/10.1016/j.patcog.2014.07.015

Foggia, P., Percannella, G., & Vento, M. (2014). Graph matching and learning in pattern recognition in the last 10 years. International Journal of Pattern Recognition and Artificial Intelligence, 28(1), 1450001. https://doi.org/10.1142/S0218001414500013

Gamble, L., Ravela, S., & McGarigal, K. (2008). Multi-scale features for identifying individuals in large biological databases: An application of pattern recognition technology to the marbled salamander Ambystoma opacum. Journal of Applied Ecology, 45(1), 170–180. https://doi.org/10.1111/j.1365-2664.2007.01368.x

Gaüzère, B., Brun, L., & Villemin, D. (2012). Two new graphs kernels in chemoinformatics. Pattern Recognition Letters. https://doi.org/10.1016/j.patrec.2012.03.020

Halloran, K. M., Murdoch, J. D., & Becker, M. S. (2015). Applying computer-aided photo-identification to messy datasets: A case study of Thornicroft’s giraffe (Giraffa camelopardalis thornicrofti). African Journal of Ecology, 53(2), 147–155. https://doi.org/10.1111/aje.12145

Huson, D. H., & Bryant, D. (2006). Application of phylogenetic networks in evolutionary studies. Molecular Biology and Evolution. https://doi.org/10.1093/molbev/msj030

Jean, C., Ciccione, S., Talma, E., Ballorain, K., & Bourjea, J. (2010). Photo-identification method for green and hawksbill turtles - First results from Reunion. Indian Ocean Turtle Newsletter, (11), 8–13.

Justice, D., & Hero, A. (2006). A binary linear programming formulation of the graph edit distance. IEEE Transactions on Pattern Analysis and Machine Intelligence. https://doi.org/10.1109/TPAMI.2006.152

Kisku, D. R., Rattani, A., Grosso, E., & Tistarelli, M. (2007). Face identification by SIFT-based complete graph topology. In 2007 IEEE Workshop on Automatic Identification Advanced Technologies - Proceedings (pp. 63–68). IEEE. https://doi.org/10.1109/AUTOID.2007.380594

Kondor, R., & Pan, H. (2016). The Multiscale Laplacian Graph Kernel. Advances in Neural Information Processing Systems, 2990–2998.

Leonardis, A., Bischof, H., Pinz, A., Bay, H., Tuytelaars, T., & Van Gool, L. (2006). Computer Vision –ECCV 2006 SURF: Speeded Up Robust Features. Computer Vision –ECCV 2006, 3951, 404-417–417. https://doi.org/10.1007/11744023

Lusseau, D. (2003). The emergent properties of a dolphin social network. Proceedings of the Royal Society B: Biological Sciences. https://doi.org/10.1098/rsbl.2003.0057

Martin-Smith, K. M. (2011). Photo-identification of individual weedy seadragons Phyllopteryx taeniolatus and its application in estimating population dynamics. Journal of Fish Biology, 78(6), 1757–1768. https://doi.org/10.1111/j.1095-8649.2011.02966.x

Miththpala, S., Seidensticker, J., Phillips, L. G., Fernando, S. B. U., & Smallwood, J. A. (1989). Identification of individual leopards (Panthera pardus kotiya) using spot pattern variation. Journal of Zoology, 218(4), 527–536. https://doi.org/10.1111/j.1469-7998.1989.tb04996.x

Myronenko, A., & Song, X. (2010). Point set registration: Coherent point drifts. IEEE Transactions on Pattern Analysis and Machine Intelligence, 32(12), 2262–2275. https://doi.org/10.1109/TPAMI.2010.46

Papandreou, G., Chen, L.-C., Murphy, K. P., & Yuille, A. L. (2015). Weakly-and Semi-Supervised Learning of a Deep Convolutional Network for Semantic Image Segmentation. In 2015 IEEE International Conference on Computer Vision (ICCV) (Vol. 2015 Inter, pp. 1742–1750). IEEE. https://doi.org/10.1109/ICCV.2015.203

Reisser, J., Proietti, M., Kinas, P., & Sazima, I. (2008). Photographic identification of sea turtles: Method description and validation, with an estimation of tag loss. Endangered Species Research, 5(1), 73–82. https://doi.org/10.3354/esr00113

Riesen, K., & Bunke, H. (2009). Approximate graph edit distance computation by means of bipartite graph matching. Image and Vision Computing, 27(7), 950–959. https://doi.org/10.1016/j.imavis.2008.04.004

Rublee, E., Rabaud, V., Konolige, K., & Bradski, G. (2011). ORB: An efficient alternative to SIFT or SURF. In Proceedings of the IEEE International Conference on Computer Vision (pp. 2564–2571). IEEE. https://doi.org/10.1109/ICCV.2011.6126544

Sanfeliu, A., Sanfeliu, A., & Fu, K. S. (1983). A Distance Measure Between Attributed Relational Graphs for Pattern Recognition. IEEE Transactions on Systems, Man and Cybernetics. https://doi.org/10.1109/TSMC.1983.6313167

Uetz, P., Glot, L., Cagney, G., Mansfield, T. A., Judson, R. S., Knight, J. R., … Rothberg, J. M. (2000). A comprehensive analysis of protein-protein interactions in Saccharomyces cerevisiae. Nature, 403(6770). https://doi.org/10.1038/35001009

Wilson, R. C., & Zhu, P. (2008). A study of graph spectra for comparing graphs and trees. Pattern Recognition. https://doi.org/10.1016/j.patcog.2008.03.011

Yuille, A. L., & Grzywacz, N. M. (1988). Motion coherence theory. In [1988 Proceedings] Second International Conference on Computer Vision (pp. 344–353). IEEE. https://doi.org/10.1109/CCV.1988.590011

